# A Novel ILP Framework to Identify Compensatory Pathways in Genetic Interaction Networks with GIDEON

**DOI:** 10.64898/2026.03.29.715009

**Authors:** Jocelyn J. Garcia, Kevin M. Yu, Catherine H. Freudenreich, Lenore J. Cowen

## Abstract

In Baker’s yeast, there exists a comprehensive collection of pairwise epistasis experiments that, for nearly every pair of non-essential genes, measures the growth of the double-knockout strain as compared to its component single knockouts. This data can be represented as a weighted signed graph termed the genetic interaction network, and we introduce a new ILP-based method named GIDEON to search for a diverse collection of Between-Pathway Models (BPMs) in this network, where BPMs are a graph motif signature that indicates potential compensatory pathways in the genetic interaction network. With both an improved distribution-informed edge weighting scheme and an improved ILP method, GIDEON produces BPM collections that are substantially larger and with better functional enrichment compared to previous methods. We find some interesting new BPM gene sets including one with potential insights into antifungal drug targets through ties between ergosterol and aromatic amino acid biosynthesis.

## 1 Introduction

Comparing the phenotype of a double gene knockout to its component viable single gene knockouts has long been known to reveal information about genes that can compensate functionally for one another; looking at this for individual gene pairs is termed the study of *epistasis*. If pairwise epistasis data is available for the growth phenotype of an organism on a genome-wide scale, this data can be represented as a signed, edge-weighted network, where the nodes are the non-essential genes and a negative (or positive) weighted edge is placed between two genes to represent how surprised one is by the sickness (or wellness) of the double deletion mutant compared to the fitness of its component single deletion mutants. Particular graph motifs in this network, termed the *genetic interaction network* have been shown to indicate higher-level epistasis, i.e. pairs of *sets* of genes, often called *pathways* that can compensate functionally for each other. Beginning with the seminal work of Kelley and Ideker [1] (more recent work reviewed in Section 2.2), a mathematical toolbox to computationally analyze the genetic interaction network has been slowly being built. However the mathematical machinery is definitely not as mature as the machinery that has been developed to analyze the protein physical interaction network (an unsigned network where nodes represent proteins and edges represent proteins for which there is experimental evidence that they bind) [2].

The best-studied genetic interaction network is currently that in Baker’s yeast (*S. cerevisiae*), where high-throughput experiments have made available measures of the fitness of single-deletion and double-deletion knockout strains, comprising nearly all non-essential yeast genes [3]. The graph motif that indicates compensatory pathways that we seek to extract many and diverse instances of is called the *Between Pathway Model*, or BPM, a variant of the original Between Pathway Model defined in the original Kelley and Ideker paper [1] (where Kelley and Ideker were instead looking at older experimental datasets [4, 5] that gave an unweighted network, where an edge between two genes indicated a *synthetic lethality* relationship, i.e. that the double knockout was completely inviable). Again, our formal BPM definition is motivated and reviewed in Section 2.1, with older work reviewed in Section 2.2.

We introduce GIDEON (Genetic Interaction-Driven Extraction of Optimal Networks), a new Integer Linear Program (ILP) based method to return a diverse collection of *Between Pathway Models* from genetic interaction networks. We compare GIDEON both to the older LocalCut method of Leiserson et al [6] and to a recent ILP method of Liany et al [7]. We find that GIDEON produces a collection of diverse BPMs whose coverage is orders of magnitude larger than previous methods, while at the same time, improving rates of functional enrichment. The gain in GIDEON comes from two main ideas: 1) a novel way to set up the ILP to make sure every node is considered for inclusion in some BPM, and 2) a better weighting scheme for capturing epistasis on the individual edges. Both methodological advancements are explained in Section 4.

## 2 Background

### 2.1 Between-Pathway Models

In genetic interaction networks, the main task is to search for *Between Pathway Models*, or BPMs. BPMs are paired subsets of genes termed *pathways* such that the sum of the weights of the edges between pairs of nodes that span across the two pathways minus the sum of the edge weights between pairs of nodes both of which lie in the same pathway is maximally negative. Figure 1 gives an example of what a typical BPM will look like: it has mostly negative weight edges (dotted in brown) going between the two pathways, and positive weight edges (solid in blue) within each pathway. If these are two pathways that can compensate for each other, but one or the other pathway is essential, then this is a typical pattern of edge weights will look like: the negative edges across represent the fact that knocking out one gene from each pathway makes the organism very sick; while knocking out two genes from the same pathway (positive edges within) results in an organism that is not as sick, since the undamaged pathway can compensate. Note that there will not, in general be observable epistasis between all genes on opposite pathways in a BPM, since there may be lower-level redundancy as well: in Figure 1, nodes *F* and *G* are intended to be paralogs that can compensate for each other, so knocking out one of them does not inactivate the righthand pathway that contains them, and so they do not have negative edges to the lefthand pathway.

**Figure 1:**
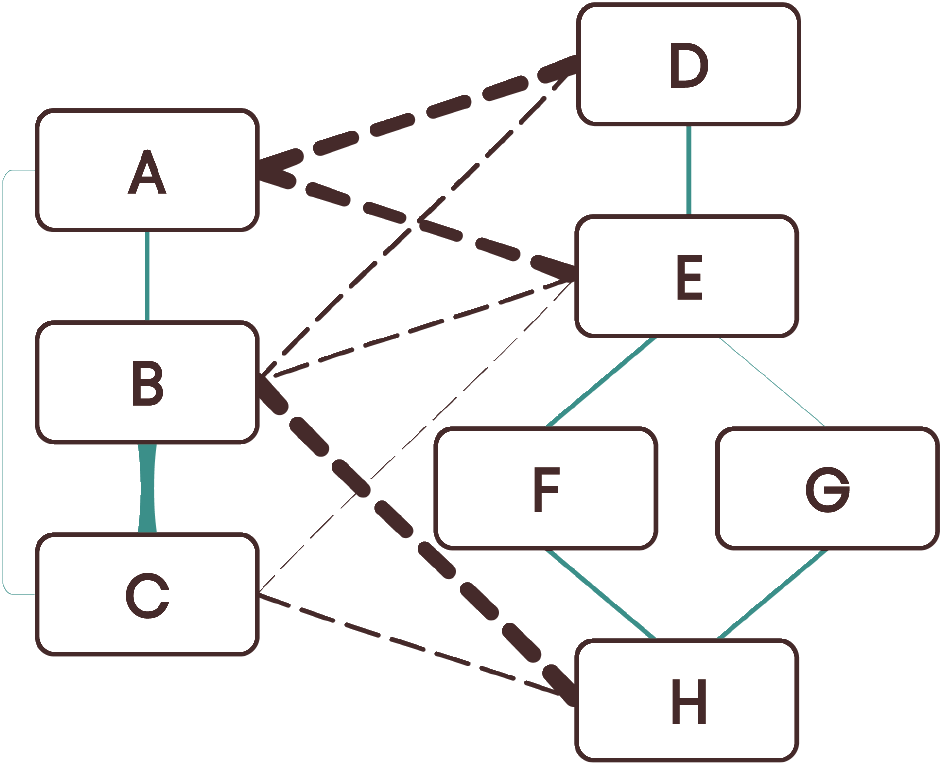
An abstract representation of the graph structure of a typical BPM. Nodes *A, B, C* participate in one biological pathway, and nodes *D, E, F, G, H* participate in a second, compensatory pathway. Solid edges represent positive interactions: knocking out two genes from the same pathway is less sick that expected because the other pathway can compensate (edge thickness represents strength of the interaction). On the other hand, dotted edges represent negative interactions: knocking out one node from each pathway, when it inactivates both pathways, makes the yeast sicker than expected. Note that nodes *F* and *G* are intended to represent paralogs that give a lower level of compensation on a single gene level, so knocking out one of them does not inactivate the righthand pathway. We would predict a triple knockout of a gene from the lefthand pathway and *both F* and *G* would be very sick, but the available data doesn’t include (most) triple knockouts.

In general, a BPM is given by naming two *modules* of genes *A* ∪ *B*, and a single number that captures the goodness of the BPM (indicating its likelihood to represent a pair of compensatory pathways) is

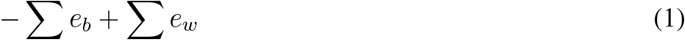

where *e*_*b*_ represent edges with one endpoint in each of *A* and *B*, and *e*_*w*_ represents edges with both endpoints in *A* or both endpoints of *B*. We seek a high-scoring set of BPMs that covers a diverse set of pathways across the entire genetic interaction network.

### 2.2 Prior Work

As mentioned above, the BPM graph motif was first defined by Kelley and Ideker in a different network setting [1]: their BPMs were defined in a network where the genetic interaction edges were unweighted negative edges (indicating synthetic lethality–the extreme case that the double knockout was inviable) and they put traditional PPI edges in place of positive genetic interaction edges to support membership in the same pathway. Subsequent work on the unweighted genetic interaction networks using principally the datasets from [4, 8] appeared in [9, 10, 11]

In the same weighted genetic interaction network setting studied here, early data came from [12, 13, 14] and early algorithms came from [15, 6], where we compare to the LocalCut from [16, 6] on the more modern comprehensive dataset of [3] below. More recently, an ILP formulation for searching for BPMs was introduced by Liany et al [7]. We observe that finding BPMs is highly similar to the problem of finding dense bipartite subgraphs of a graph, which is NP-hard. Thus one expects it to be difficult to to find good methods to search for collections of BPMs, and so an ILP formulation that can be solved heuristically in practice seems to be a good strategy. However, as we will show below, our ILP formulation has several advantages over the ILP defined by Liany et al. [7]

When comparing different BPM-finding algorithms, the goal is to find as large and diverse a collection of BPMs as possible. Typically, the diversity is enforced by giving a minimal *Jaccard Index* and requiring that two BPMs returned by the same collection have bounded overlap (in Jaccard measure) (See Section 4.3 for formal definitions). It doesn’t really matter in practice exactly how this threshold is set as long as it is bounded away from 0 and 1: however, it should be set the same in all compared methods for consistency. Thus we set the threshold here at.66 because that’s what has been done in every previous paper [15, 6, 17]. Then, we can compare collections of BPMs filtered by this allowed overlap percentage with two different measures of goodness, as is also standard in the field: 1) sheer size of the collection, or number of BPMs returned (enforcing the Jaccard overlap constraint) and 2) functional enrichment for the component pathways of the BPMs we discover. We next briefly review the two prior BPM methods we compare against GIDEON.

#### LocalCut

LocalCut, described in [16, 6], balances the tension between local and global methods of BPM discovery by greedily performing many partitions of the interaction network that come from global maximal cuts. Each gene is then used to form the modules of a BPM by including other genes that co-locate to either the same or opposite partition in at least *c*% of the partitions.

#### Liany-ILP

Liany et al. [7] introduced a method for discovering BPMs, the first ILP formulation to our knowledge. More specifically, we compare against their “Top-K Maximum Weight Bipartite Subgraph” method, but we will call it Liany-ILP here. Their approach uses an ILP to sequentially identify the BPM that maximizes the weight of negative interactions across modules, ignoring any interactions within BPM modules. Once a BPM is identified, the edges of said BPM are removed from the network and the ILP is run again on the subnetwork. Given time limitations on solving a large ILP, the user specifies a time limit for the ILP to search for a BPM, and the algorithm halts when all edges of the network have been removed. The only user-specified constraints are the number of BPMs (K) and their sizes.

## 3 Approach

We introduce GIDEON, a new ILP-based method to return such a collection of BPMs, that we show returns a much more comprehensive set of diverse enriched BPMS on the yeast genetic interaction dataset of [3]. The gains of GIDEON come from two main things: first, we improve the weighting scheme of Yu and Cowen [18] to better model epistasis weights of the double knockouts (see Section 4.5). Second, we introduce a new ILP formulation for the problem that uses a clever trick to allow the ILP to discover diverse pathways, where most obvious ways to define a BPM search as an optimization problem tend to converge on a single “best” BPM over and over again. With our formulation, unlike that of Liany et al. [7], we need not require interactions from previously discovered BPMs to be permanently removed from consideration in order to discover new gene sets. We find that by all previous metrics, number of BPMs, number of BPMs enriched for known function, that GIDEON substantially outperforms prior methods for BPM discovery, from the LocalCut method of [16, 6] to the recent ILP formulation of Liany et al. [7]

## 4 Methods

When constructing GIDEON, we aim for two goals: individual BPMs are filled with biologically coherent sets of genes and the BPM outputs are diverse. We meet the second goal using the approach of previous methods, by building a larger set of outputs and pruning BPMs based on their Jaccard similarity to other BPMs in the final output set. For constructing BPMs, we actually solve a different ILP centered around each gene in turn, echoing the gene-centered approach of LocalCut. Finally, our ILP is constructed on a network which is both reweighted, to better capture epistasis, and sparsified, removing low weight edges which helps make the ILP computationally tractable (see Sections 4.5 and 4.6 below).

### 4.1 Constructing the ILP

GIDEON solves the following ILP for every nonessential gene *g*_*c*_, where *E*_*c*_ represents the set of edges adjacent to *g*_*c*_, *E*_*g*_ represents the set of edges adjacent to an arbitrary gene in the network *g*, and *w*_*e*_ represents the *squared* weight of the edge *e* in the genetic interaction network (retaining the positive or negative):

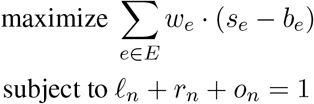

Gene-Centered Constraints:

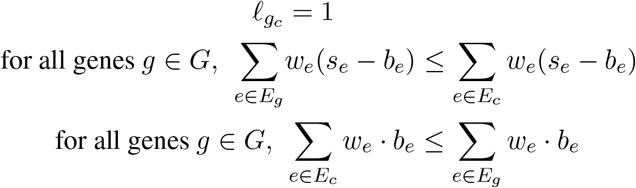

Edge-Node Constraints:

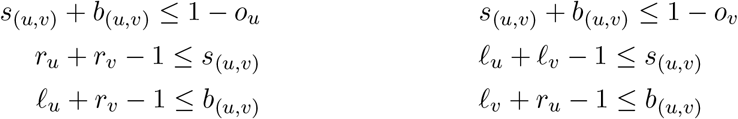

Size Constraints:

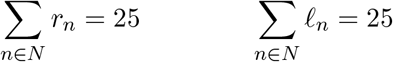

Node and Edge Variables:

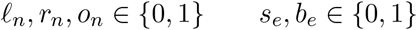

Before discussing the intuition of GIDEON, we notate the location of a node *n* within the BPM using the binary node variables such that the corresponding variable is 1 when the node is in the *left* module, *right* module, or *outside* of the BPM, meaning that it is not included in either gene set. The first constraint then ensures each node is either in exactly one BPM module or considered “outside” of the BPM. To calculate the objective function based on edges between and within the same module, the binary edge variables represent whether a given edge *e* = (*u, v*) is within the *same* module or *between* modules, with both variables set to 0 when an edge is not included in the BPM.

Using the above notation, the ILP portion of GIDEON consists of two key components: the constraints that ensure local optimization about a specific gene *g*_*c*_ while promoting cohesive BPM modules and the bare-bones encoding of the BPM problem into an ILP that specifies the location of genes and edges in the BPM. For each gene *g*_*c*_ in the network, we direct its associated ILP to build a BPM that maximizes an objective function that rewards negative edges between BPM modules and positive edges within BPM modules, where *w*_*e*_ represents the edge weight and thus the nature and strength of the genetic interaction between a pair of genes. Note that we square all weights, as recommended by [6, 18] and discussed in Section 4.5. This objective function is a proxy to the interaction weight from [6] used in Section 4.3, which includes a nonlinear aspect to account for BPM size. Given that large BPMs naturally maximize this function, we later introduce a size constraint to the ILP accompanied by BPM trimming for removing loosely connected genes. We imbue the local flavor of LocalCut into the global nature of an ILP by ensuring that the BPM constructed by each gene’s ILP not only includes *g*_*c*_ but is constructed based on the interactions of *g*_*c*_. The gene-centered constraints ensure that *g*_*c*_ is in one of the modules, *g*_*c*_ is the primary contributor to the objective function out of all the genes included in the BPM, and that *g*_*c*_ is specifically the primary contributor to the negative interaction across the BPM modules. These constraints direct the ILP towards BPM structure centered around *g*_*c*_, keeping the ILP from superficially adding *g*_*c*_ to an unrelated strong module. *Because every BPM must have a primary contributor to the objective function, these constraints elegantly ensure that possible BPMs are not removed from the search space at the expense of centering on a given gene*.

Having centered the ILP on *g*_*c*_, note that we still must require the Edge-Node constraints in the ILP shared across all genes that impose biconditional relationships between then node and edge variables. For the edge *e* = (*u, v*), namely

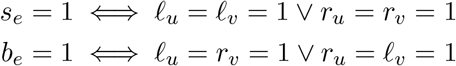

To impose these relationships, the constraints implement conditional statements for each case that, in conjunction, ensure that the biconditional relationships hold. When either endpoint is outside of the module, the first pair of constraints enforces *s*_*e*_ = *b*_*e*_ = 0 while simultaneously ensuring that an edge cannot be defined as both between and within modules (in that *s*_*e*_ + *b*_*e*_ ≤ 1). The second pair of constraints then enforces that *s*_*e*_ = 1 when *u* and *v* are in the same modules, but these constraints have no impact when that condition is not fulfilled. The third pair of constraints similarly enforces *b*_*e*_ = 1 when *u* and *v* are in opposite modules while not impacting nodes without this behavior.

A desire for BPMs of reasonable size last informs a set of natural size constraints for the ILP. In the absence of these constraints, one can imagine that a notable number of genes in the large set of nonessential genes will at least minimally improve the objective function. In addition, the computational constraints for searching for the optimal sets of genes blow up exponentially, meaning that some limit is required for the ILP to be computationally feasible. We thus hardcoded a limit on the maximum number of 25 genes for each module, where we chose 25 because it was still computationally tractable to solve the ILP and it was consistent with previous work (25 is the an upper bound on BPM size in LocalCut [6]). However, our intuition that the ILP can always find genes that weakly improve the objective function is correct: when we solve the ILP, it almost always returns 25 genes in each module. But some of these genes clearly seem like “extra” genes that are not important to the BPM, so we then trim the BPM, as described next. The trimming procedure is so effective, that we instead ask the ILP to always return 25 genes exactly in each module (rather than a maximum of 25 genes) and then trim, improving the computational complexity of its search space.

### 4.2 Trimming BPMs

Given the initial constraint for exactly 25 genes in each module, we remove weakly related genes and thus trim the BPMs. Every gene *g* with only one negative interaction across the modules or has a gene interaction weight less than 0.015, meaning that

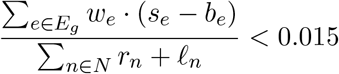

is removed. To ensure that every remaining gene maintains these criteria even after others are removed, we iteratively remove genes until none are removed. Once trimmed, BPMs with less than 3 genes in either module are removed, consistent with the minimum size constraints of [6]. We chose a cutoff of 0.015 after finding robust results within this range (Figures B.1, B.2).

### 4.3 Pruning BPMs

Consistent with [6], the trimmed set of BPMs from GIDEON are pruned by their similarity, ensuring diverse results. BPMs are first ranked by their interaction weight (without edge squaring), defined as Equation 1 divided by the BPM size. Beginning with the highest interaction weight, we add BPMs to the final result set if their Jaccard similarity is less than 0.66 with every previously added BPM, where the Jaccard similarity for a pair of BPMs (*A*_*x*_, *B*_*x*_), (*A*_*y*_, *B*_*y*_) is defined as

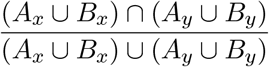

### 4.4 Final Trim

While the above excels in finding BPMs that maximize the ILP objective, i.e. in putting sets of genes with large negative weights in opposite pathways, the union of multiple BPMs also performs well under the ILP objective function. In fact, we find 26% of BPMs returned by the ILP after trimming form *>* 2 connected components in the sparse network that includes only the most significant edges (see Section 4.6) and only interactions with magnitude ≥ 0.2. This is less than the 46% such BPMs that form *>* 2 connected components by LocalCut and the 79% by Liany-ILP. Therefore we break up BPMs to produce the final GIDEON collection as follows: after removing all edges with magnitude *<* 0.2, we add all connected components of BPMs with at least 3 genes in each module to the final set of “core” BPMs, followed by the same pruning from Section 4.3 to ensure diverse sets. In Figures B.9 and B.10, we also compare our BPMs to LocalCut and Liany-ILP with the same Final Trim applied to their BPM collections and find that GIDEON further distances itself in BPM quantity and enrichment.

### 4.5 Weighting the Edges

While the ILP formulation is allowing us to optimize the best collection of weighted BPMs according to the objective function, the BPMs we find will only be as good as the epistasis values that are captured on the edge weights. There are several plausible null-models and thus equations that can be used to compute the expected weight of a double knockout given the growth rates of its component single knockouts: Mani et al [19] explored empirically what produced the best edge weights to detect epistasis on a single edge basis. Cowen and Yu [18] revisited this question in the context of looking for entire pathways that participated in a BPM, not just on an edge by edge basis. In particular, they showed that the suggestion of [16] to *square* edge weights (but retaining the original sign), pulling the smallest weights toward zero and increasing the relative magnitude of the largest weights, improved performance of the LocalCut algorithm because it served to denoise the network on the pathway level. We also find that sparsifying the network by throwing out low-magnitude edges improves the performance of LocalCut (see Section 4.6 and Figures B.3,B.4).

GIDEON also benefits from squaring edge weights and network sparsification, but makes a conceptual advance in weighting schemes beyond previous work. Instead of computing the edge weight as a function of just two genes, GIDEON examines the entire distribution of a given gene’s double knockouts in order to have a more robust sense of which double knockouts are outliers. By doing so, it more precisely characterizes each gene’s typical response as a component in a double knockout and therefore gains a targeted understanding of when a double knockout deviates from that norm. We note that a similar weighting scheme for double knockouts was first proposed by [20] as a minor adaptation of [21]. We show that this improved way of assigning edge weights, which we will refer to as “Distribution Informed” (DI) edge weights, also improves the performance of the competitor methods (Figures 3 and B.5) we test against. Thus, it is a stand-alone contribution of this paper (in addition to the ILP).

To introduce our DI weighting scheme for the gene interaction network, we formalize the notion of a double knockout marginal distribution. For a gene *a*, the *marginal distribution* is the single mutant fitness *S*_*b*_ of the second component gene *b* in the double knockout relative to the double mutant fitness of the double knockout *D*_*a,b*_ for all genes *b*. Preliminary analysis of the marginal distributions demonstrates strong linear relationships between the single and double mutant fitnesses, exhibiting a tie to the success of the multiplicative model because, when conditioning for a given gene, a multiplicative model will exhibit this behavior. For our proposed weighting scheme, we use the residual of a double knockout’s true fitness from predicted by linear regression on the marginal as a measure of epistasis. Because there are two such predictions for the marginals on both component genes in a double knockout, we define the edge weight as follows:

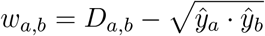

where *ŷ*_*a*_, *ŷ*_*b*_ represent double mutant fitness predictions from the marginal distributions of *a* and *b* respectively and *D*_*a,b*_ represents the true double knockout fitness. Usage of the geometric mean reflects the previous success of a multiplicative model and the use of linear regression allows the residual standard deviation *s*_*res*_ to act as a proxy for edge confidence, quantifying the predictability of gene behavior as a double knock-out component.

Note that a suppressor mutation in a small number of strains led to the artifact in Figure 2D, which we account for by constructing separate marginals for the gene acting as the *array* or *query* (further details found in Section A.2).

**Figure 2:**
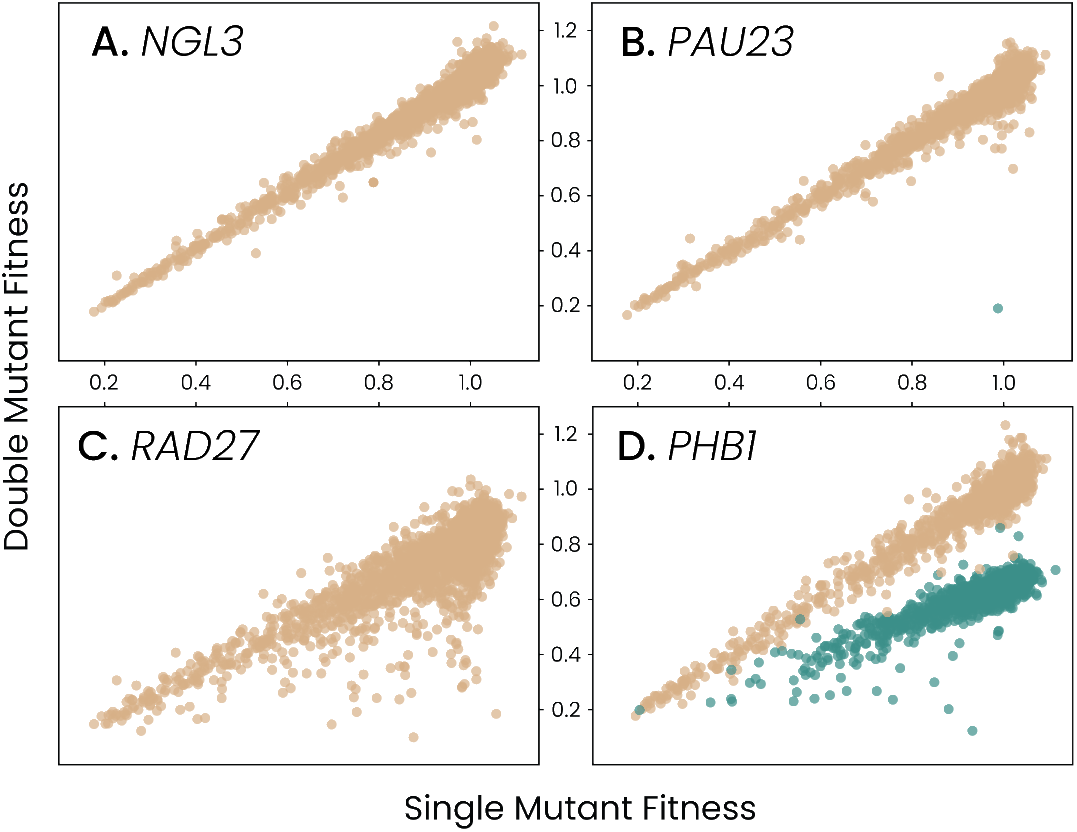
The marginal distributions for each gene tend to fall in the above four categories. Many genes demonstrate the highly linear behavior of *NGL3* in (A), with a number of genes also including a few outliers of synthetic sickness interactions shown by *PAU23* in (B). Some genes exhibit less predictable behavior as double knockout components such as *RAD27* in (C), resulting in higher variance in the model used for our DI weighting scheme and therefore a stricter requirement for a double knockout to be seen as surprising. Last, a very small number (37) of genes demonstrated the split trajectory shown by *PHB1* in (D), with the colors representing whether *PHB1* was the *array* or *query* gene in the synthetic genetic array data (an experimental artifact from a suppressor mutation in the strain).

### 4.6 Network sparsification

We filter the genetic interaction network by the minimum standard deviations from predicted in the marginal distributions of each component gene, removing all interactions with *s*_*res*_ ≤ 2. Upon filtering, three genes (*PEP3, BUD22, VPS16*) have no remaining interactions. LocalCut performance on successive edge filterings showed that this strict setting identified the largest number of BPMs from the network with competitive enrichment statistics (Figure B.3). By preserving only 2% of edges, we also observe a significant reduction in the quantity of variables required for GIDEON, lessening the expense of solving the ILP. To more carefully explore whether GIDEON’s gains come from the ILP or the new weighting schemes, we tested all of GIDEON, LocalCut and Liany-ILP using different weighting schemes in Figure 3, spefically using the original weights from [3] with their suggested “stringent” filtering. Results from their “intermediate” filtering as well as a magnitude-based filtering of the multiplicative weighting scheme also appear Figure B.5.

**Figure 3:**
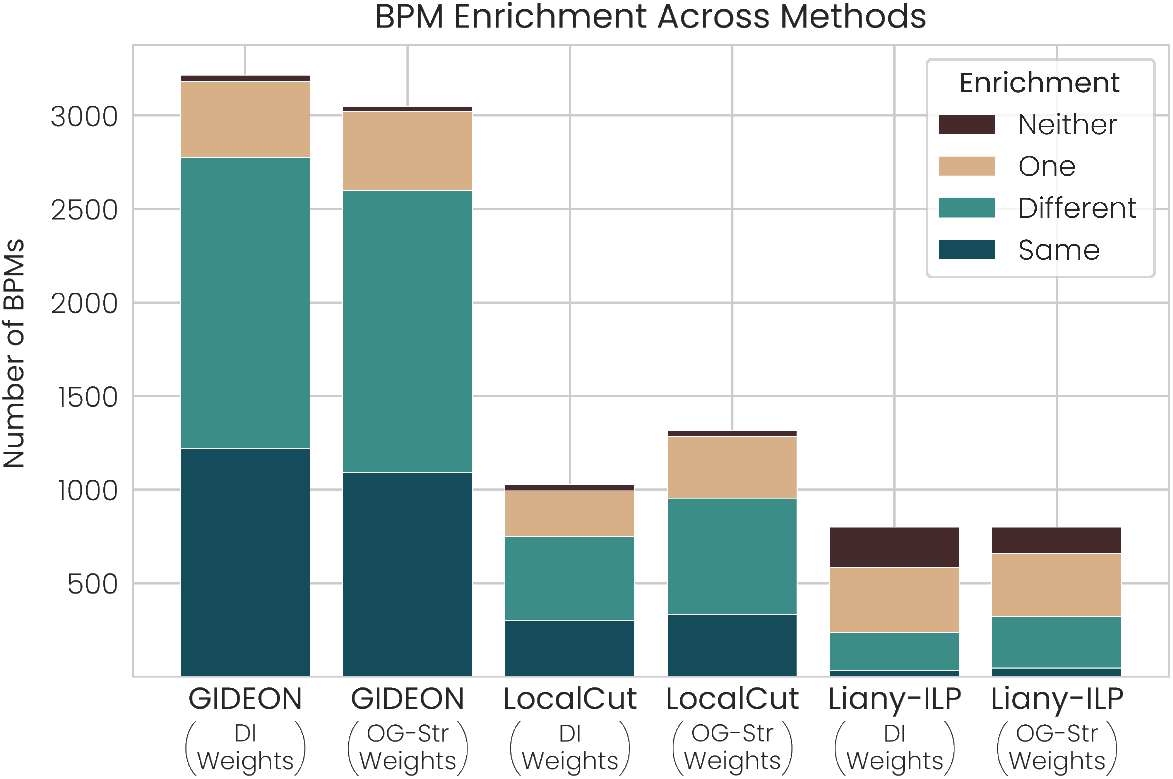
GIDEON with our new DI weights shows massive improvements both in number of BPMs found and in their enrichment. GIDEON with the stringent filtering of [3] weights is second best. Previous LocalCut produces substantially fewer BPMs, even using our new DI weighting scheme and that of [3]

### 4.7 Pathway Enrichment Analysis

To measure BPM coherence, we compare the enriched terms of each module, categorizing BPMs by whether their modules are enriched for the same term, different terms, or if only one or neither of the modules are enriched. Modules were evaluated against the GO, KEGG, WikiPathway, and Human Genotype Ontology gene sets using the GProfiler package [22] with the Benjamini-Hochberg [23] adjustment for multiple testing. Similar to [18], and keeping with general practice in the field, we only consider enrichment by GO terms that label at most 500 different genes, in order to exclude the most general levels of the GO hierarchy.

### 4.8 Solving the ILP

To solve the ILP, our implementation exclusively uses the No Relaxation (NoRel) heuristic in the Gurobi solver [24] oriented toward problems with challenging root LPs or a large number of binary decision variables. Given the nature of our ILP, using only the NoRel heuristic proved notably more efficient than techniques that rely on solving the root LP, quickly locating BPMs with stronger objectives by solving sub-ILPs in parallel and piecing together their solutions. The heuristic can only be controlled by time/computation limits, and we found that 300 work units produced coherent BPMs. Note that the full ILP script required an average of 8.5 minutes and *<* 5 GB of memory per gene on an Intel(R) Xeon(R) Gold 6438M using up to 8 threads.

## 5 Results

GIDEON with our DI weighting scheme identifies nearly three times more BPMs than its best performing competitor, being LocalCut with the stringent filtering from [3] (referred to as OG-Str). GIDEON consistently outstripped previous methods when the same network is used and performed best with the DI weighting scheme on the metrics of BPM quantity, module size, and enrichment when compared OG-Str. In this setting, GIDEON identifies 3,215 BPMs, whereas LocalCut and Liany-ILP only identify 1,027 and 750 BPMs respectively. On the measure of BPM enrichment, Liany-ILP performed particularly poorly and identified 33 BPMs enriched for the same function, while LocalCut identified 301 and GIDEON identified 1,220.

Though the OG-Str weights from [3] slightly outperformed our DI weights for LocalCut and Liany-ILP (though not for GIDEON), we remark that computing these weights require additional experiments and knowledge of array positions and batch effects, which will not be available for every dataset. Computing DI weights in contrast, requires only single and double mutant fitnesses. LocalCut tended to perform better on filtered networks, but comparisons on the filtered version of the multiplicative weights then also demonstrate that gains provided by our DI weighting scheme are not simply from using a smaller network either (Figure B.3). A full comparison of all the different methods with all the weighting schemes appears in Section A.3 and Figures B.5 and B.6.

Similar to [18], we also compared LocalCut performance on our DI weighting scheme to previous logarithmic, multiplicative, and minimum weighting schemes and found that the DI weighting scheme outperformed the others, with one filtering of the weights from [3] providing competitive results (Figures B.3, B.4). The distinction between these results and [18] is that we ran LocalCut on the complete interaction network rather than a subgraph.

For the competitor methods, we ran LocalCut using *c* = 70 and squared the edges following a hyperparameter search (Figures B.3, B.4). For Liany-ILP, we used the GIDEON settings of up to 8 threads with a work limit of 300 units and used *K* = 750 after observing poor BPM enrichment (Figure B.8). Consistent with LocalCut and GIDEON, Liany-ILP had size constraints between 3 and 25, and we pruned output to ensure consistent BPM diversity measures. While GIDEON and LocalCut BPMs had average modules sizes of approximately 9 genes in Figure 3, Liany-ILP always included the weakly improving genes that our trimming procedure addresses and averaged 25 genes per module.

## 5.1 Distinct Overlapping BPMs

Two BPMs outputted by GIDEON in Figure 4 contain genes involved with homologous recombination, with unique genes in BPM A emphasizing replication fork processing and in BPM B the nuclear pore. Together, both BPMs give a broader landscape of potential genetic pathway interactions. We observe that, based on the way they formulate the ILP, [7] will never find both these BPMs in its BPM collection, since the four shared edges would be removed from the network by [7] after discovering one.

**Figure 4:**
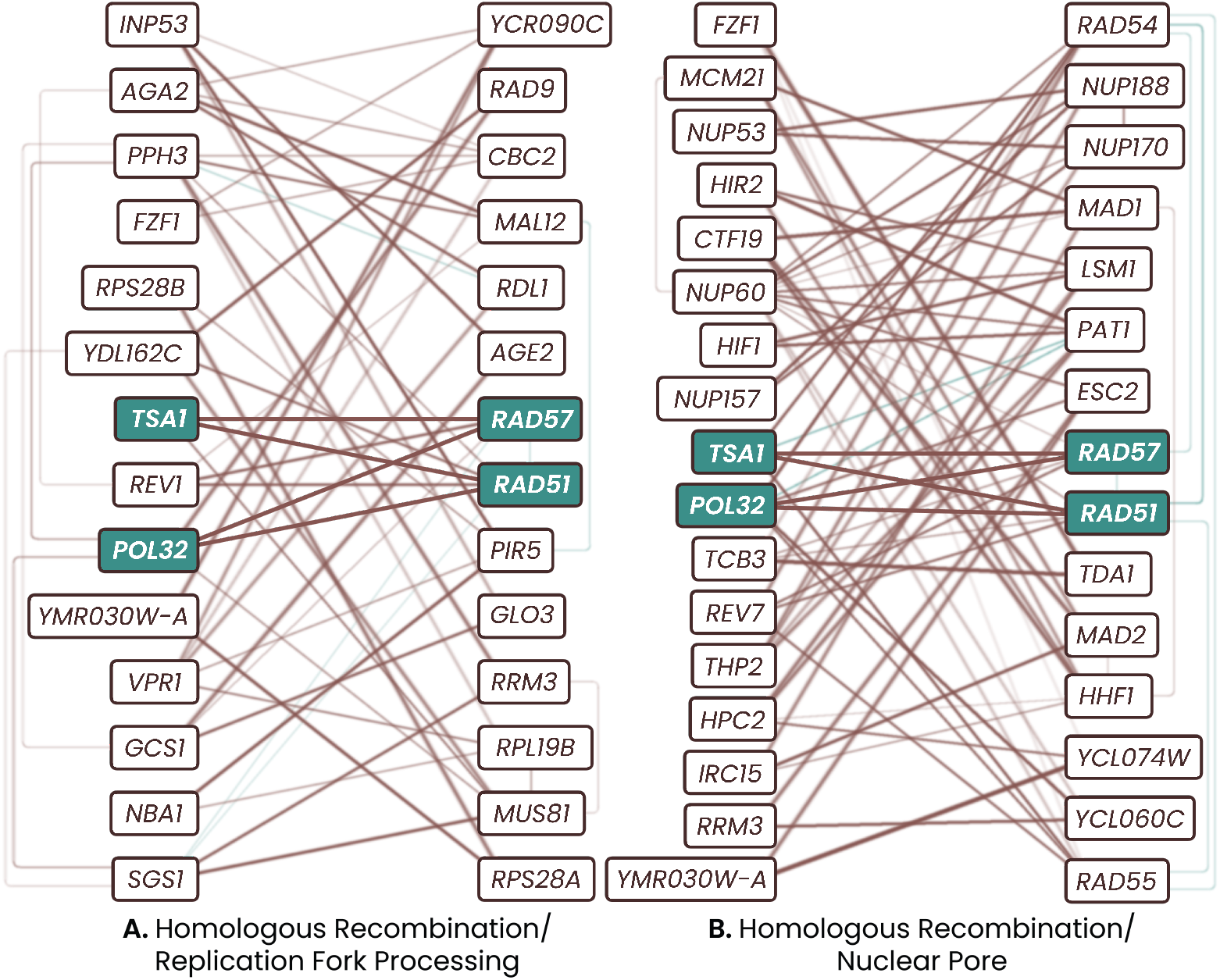
GIDEON allows interactions to be included in multiple BPMs, with the shared interactions and their component genes highlighted above. Brown edges are negative interactions and teal edges are positive interactions.

### 5.2 Ergosterol & Aromatic Amino Acid Biosynthesis

We highlight an interesting BPM in Figure 5 centered on TKL1, whose pathways are enriched for distinct functions: the left for aromatic amino acid biosynthesis and the right for ergosterol biosynthesis. Ergosterol is a critical fungal cell membrane steroid for stress adaptation and an antifungal drug target [25]. Multiple genes in the right module are either directly involved in to ergosterol biosynthesis or demonstrate ties to amino acid transport. *ERG3* and *ERG6* are in the ergosterol biosynthesis pathway, with *ERG6* specifically suggested as an antifungal target [25]. *VBA5* and *BAP2* are each plasma membrane proteins involved in amino acid uptake, with *STP1* being the transcription factor of *BAP2* [26].

**Figure 5:**
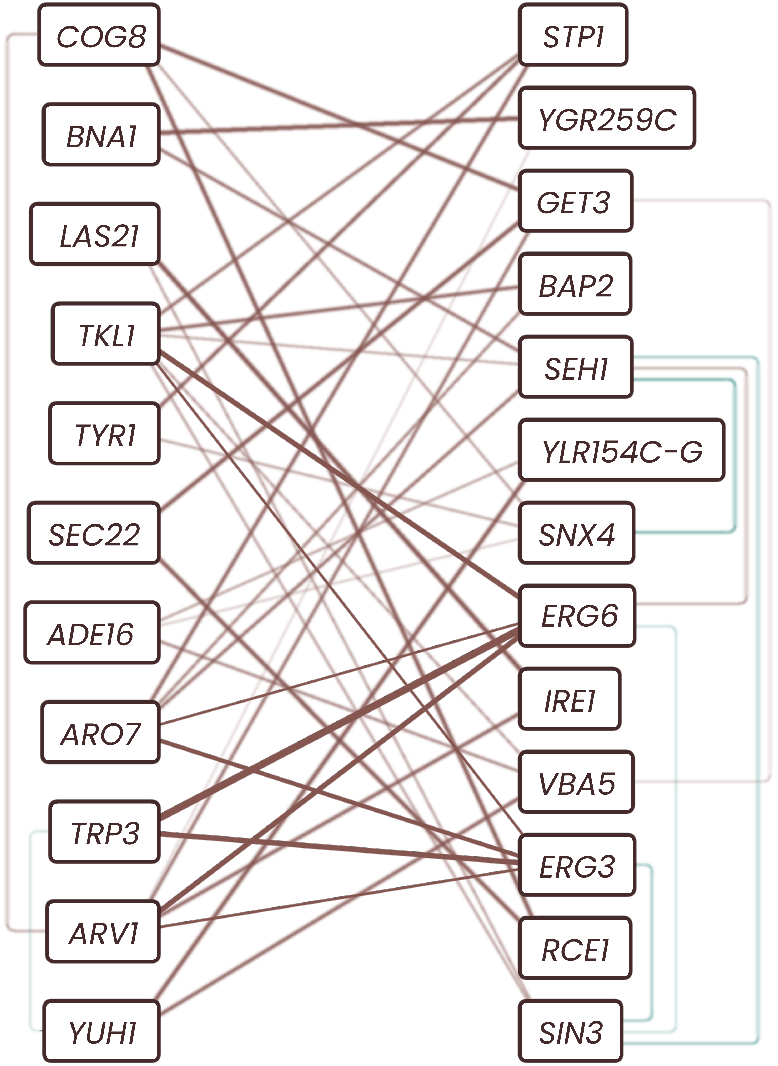
A BPM highlighting the tie between ergosterol and aromatic amino acid biosynthesis: negative interactions in brown primarily across the modules and positive interactions in teal, with strong interactions between *ERG3, ERG6* and *TRP3, ARV1, ARO7, TKL1*.

The left module boasts three shikimate pathway enzymes (*TRP3, ARO7, TYR1*), a key piece of aromatic amino acid synthesis that has been described as an attractive source of antifungal drug targets [27]. Though YGR259C is a dubious open reading frame, it almost completely overlaps with *TNA1* and thus its interaction profile likely resembles *TNA1* [28]. The role of *TNA1* in related processes further suggests this resemblance, with *TNA1* and *BNA1* each involved in NAD^+^ biosynthesis from the aromatic amino acid tryptophan [29]. *TKL1* is also required for aromatic amino acid biosynthesis [30], while *ARV1* relates to ergosterol through involvement in sterol transport and exogenous sterol uptake [25].

Inclusion of amino acid uptake genes with *ERG3* and *ERG6* across from amino acid biosynthesis suggest potential compensatory relationships between the two processes. Previous work has suggested inhibitors of aromatic amino acid biosynthesis as antifungal drugs [27] with tryptophan biosynthesis noted as particularly promising [31]. The relationships identified above could thus suggest potential insight into future antifungal drugs.

### 5.3 Frequently Included Genes

Without implicit limitation on the number of BPMs that can include a given gene or interaction, GIDEON allows for the ubiquity of genes involved in general stress response, detailed in Table 1. The most common genes in GIDEON BPMs tend to specifically play roles in unfolded protein response, Golgi transport, and DNA damage response.

**Table 1.**
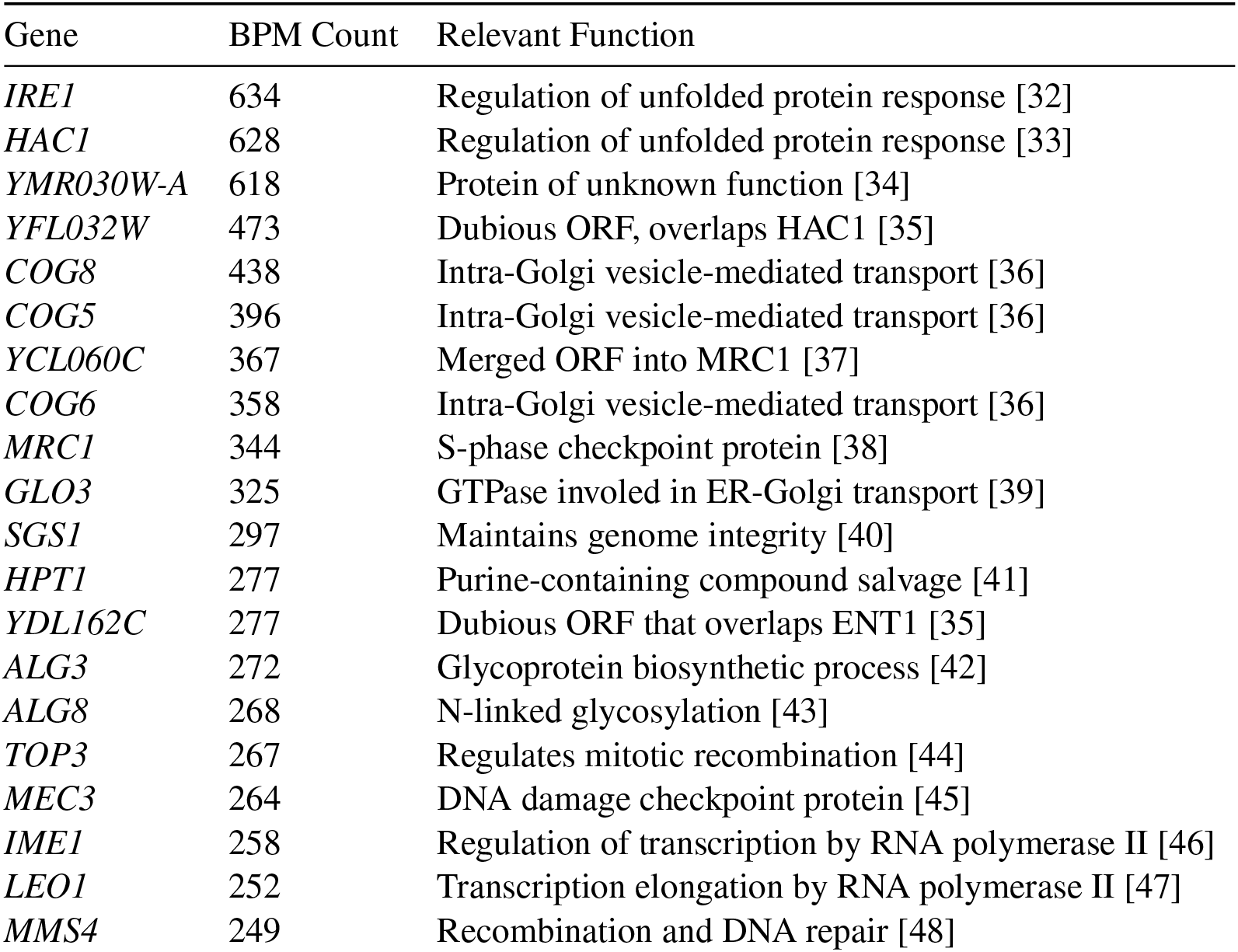
The twenty most common genes across GIDEON BPMs using our DI weighting scheme, sorted by their frequency.

## 6 Discussion

We have presented GIDEON, a novel ILP formulation coupled with a new edge-weighting scheme for the problem of searching for putative compensatory pathways in genetic interaction networks that returns a much larger and more functionally enriched collection of such BPM motifs in the yeast genetic interaction network. Beyond yeast, we are interested in adapting and generalizing GIDEON to search for compensatory pathways that might be druggable in the treatment of human cancers. High-quality high-throughput genetic interaction data is just beginning to emerge that would allow this exploration [49]. A recent paper from the group that produced Liany-ILP [50] has initial work on predicting cancer-relevant synthetic lethal interactions and we are excited to explore this further.

## 6.1 Availability and Implementation

Code and the full set of BPMs we uncover are available at https://github.com/jocelynjgarcia/GIDEON/

## 6.2 Acknowledgments

Thanks to Carol Kumamoto for valuable insights into ergosterol function, Faith Occiti and Will White for code review, and the Tufts BCB group for helpful discussions. We thank the National Institute Of General Medical Sciences of the National Institutes of Health under Award Number R01GM163241. KMY thanks the DIAMONDS REU under NSF grant 2149871. The content is solely the responsibility of the authors and does not necessarily represent the official views of the National Institutes of Health.

## Appendix

### A Extended Methods

#### A.1 Constructing the Nonessential x Nonessential Gene Network

For some pairs of genes *a* and *b*, the nonessential double knockout data from [3] contains reciprocal double mutant fitnesses: one corresponding to the cross where *a* is the query strain and *b* is the array strain, and another where the roles are reversed. After removing interactions with missing values in either the single or double mutant fitnesses, we use the most lenient suggestion from [13] as follows:

1. Remove both reciprocal interactions when the interactions from [3] have opposite signs (say, *a* as the query strain has a positive interaction and *b* as the query strain has a negative interaction).
2. For any remaining duplicate edges, remove the interaction with the higher p-value according to [3].

After removing edges with missing values and following their suggested filtering, the resulting genetic interaction network has 4,459 genes with 6,821,060 edges.

#### A.2 Adjusting for Experimental Error in the DI-Weighting Scheme

As discussed in [3], a small number of single mutant strains were found to carry a second suppressor mutation, leading to the behavior in the marginal distribution demonstrated by *PHB1* in Figure 2. All strains marked with a suppressor mutation demonstrated different single mutant fitnesses based on whether they were in the array or query strain in the double knockout. Across the data, we identified 37 genes whose single mutant fitnesses differ between the array and query strains, and these 37 genes include all strains with suppressor mutations as well as additional genes whose strains may have carried suppressor mutations as well based on their marginal distributions. The genes with differing single mutant fitnesses, as well as the magnitude of that difference, are as follows: *LTE1* (0.05), *UPS1* (0.22), *NUP170* (0.26), *HCR1* (0.30), *ARV1* (0.31), *RSM25* (0.57), *YIL014C-A* (0.02), *NCE101* (0.01), *PEX32* (0.07), *CBF1* (0.33), *MSC1* (0.31), *THP3* (0.31). *RPL43A* (0.34), *SNF1* (0.27), *VPS53* (0.29), *REI1* (0.18), *RRP6* (0.35), *SLX8* (0.08), *MNN10* (0.26), *PHB1* (0.35), *PAC10* (0.34), *LPD1* (0.12), *DDR2* (0.01), *GTR2* (0.13), *SAE3* (0.02), *ACL4* (0.01), *RMR1* (0.14), *BUB1* (0.24), *ROM2* (0.37), *LHS1* (0.12), *BUD27* (0.13), *SWA2* (0.28), *ARC18* (0.30), *SEC22* (0.31), *MRM2* (0.31), *DCK1* (0.01), *MAC1* (0.02).

When computing the weights and the residual standard deviation *s*_*res*_ for these 37 genes in the Distribution-Informed Weighting Scheme, we constructed two separate marginals, each with their own model and *s*_*res*_, based on whether the gene was in the query or array strain in a double knockout. This allowed the model and *s*_*res*_ to more accurately reflect the gene’s typical behavior in a double knockout.

In [3], there are also a small number of genes (17) with two different array single mutant fitnesses, but these unique values differed by less than 0.025 and therefore do not present a strong enough argument to build separate marginal distributions within the array marginal distribution. That said, in the event of significant discrepancies, one could imagine generalizing the Distribution-Informed waiting scheme to build separate marginal distributions based on the array single mutant fitness, forming more homogenous groups with a more accurate measure of expected gene behavior in that environment.

We last note that the computation of residual standard deviation has a divisor of *N* − 2, so the edge-case where a model is built on less than 3 data points can cause a negative or undefined *s*_*res*_. Though this event did not occur in the data used here (and should not occur with high-throughput SGA data), when deciding which edges to filter, the implementation of GIDEON currently opts to keep any interaction based on less than 3 data points rather than attempt to compute residual standard deviation, with the argument that such few data points do not give a true sense of expected gene behavior.

#### A.3 Additional Weighting Schemes for Comparison

In addition to comparing GIDEON, LocalCut, and Liany-ILP using our DI weights and the stringent filtering of [3] weighting schemes, we also compared each method performance to the intermediate filtering of [3] as well as the weighting schemes compared by [18] from [19]. As discussed in Section 5 of the main text, LocalCut had superior performance on smaller networks and the use of filtered networks allows for manageable model sizes in GIDEON and Liany-ILP. Costanzo et. al. provided suggestions for filterings of the network described as stringent and intermediate, each offering different balances of false negatives and false positives [3].

- *Original-Stringent*: Retain all edges where *P <* 0.05 and |*ε*| *>* 0.08
- *Original-Intermediate*: Retain all edges where *P <* 0.05 and *ε >* 0.16 or *ε <* −0.12.

With our DI Weighting Scheme containing 156,428 edges, these two filterings provide an elegant comparison to networks with slightly fewer and slightly more edges, with the Original-Stringent and Original-Intermediate filterings producing networks with 97,162 and 250,081 interactions respectively. We will refer to these weighting schemes as OG-Str and OG-Int.

Beyond the filterings of the original weights, we compared LocalCut performance using the following weighting schemes discussed in [19, 18], where *D*_*a,b*_ represents the double mutant fitness of genes *a, b* and *S*_*a*_, *S*_*b*_ represent the single mutant fitnesses of genes *a, b* respectively:

- *Minimum*: *w*_*a,b*_ = *D*_*a,b*_ − min(*S*_*a*_, *S*_*b*_)
- *Logarithmic*: *w*_*a,b*_ = log_2_(*D*_*a,b*_) − (log_2_(*S*_*a*_) + log_2_(*S*_*b*_))
- *Multiplicative*: *w*_*a,b*_ = *D*_*a,b*_ − (*S*_*a*_ · *S*_*b*_)

Note that we will refer to the weighting schemes as Min-All, Log-All, and Mult-All to represent that the full network of interactions is being used, while the abbreviation “Mult” specifically refers to the multiplicative weighting scheme filtered by edge magnitude to the same size as the DI weights (156,428). We similarly refer to the DI weights on the full interaction network without the filtering by residual standard deviation as DI-All. Though [3] stated that they computed their interaction scores using the multiplicative weighting scheme, their data presents a small number of discrepancies. Of the 6,821,060 unique interactions in our network, 20 had weights of 0.0 rather than their expected weights under the multiplicative weighting scheme. A very small number of interactions (119) also had negative double mutant fitnesses, which we removed from the data only when computing the Logarithmic weighting scheme because a negative double mutant fitness produces an undefined interaction score.

### B Additional Figures

**Figure B.1:**
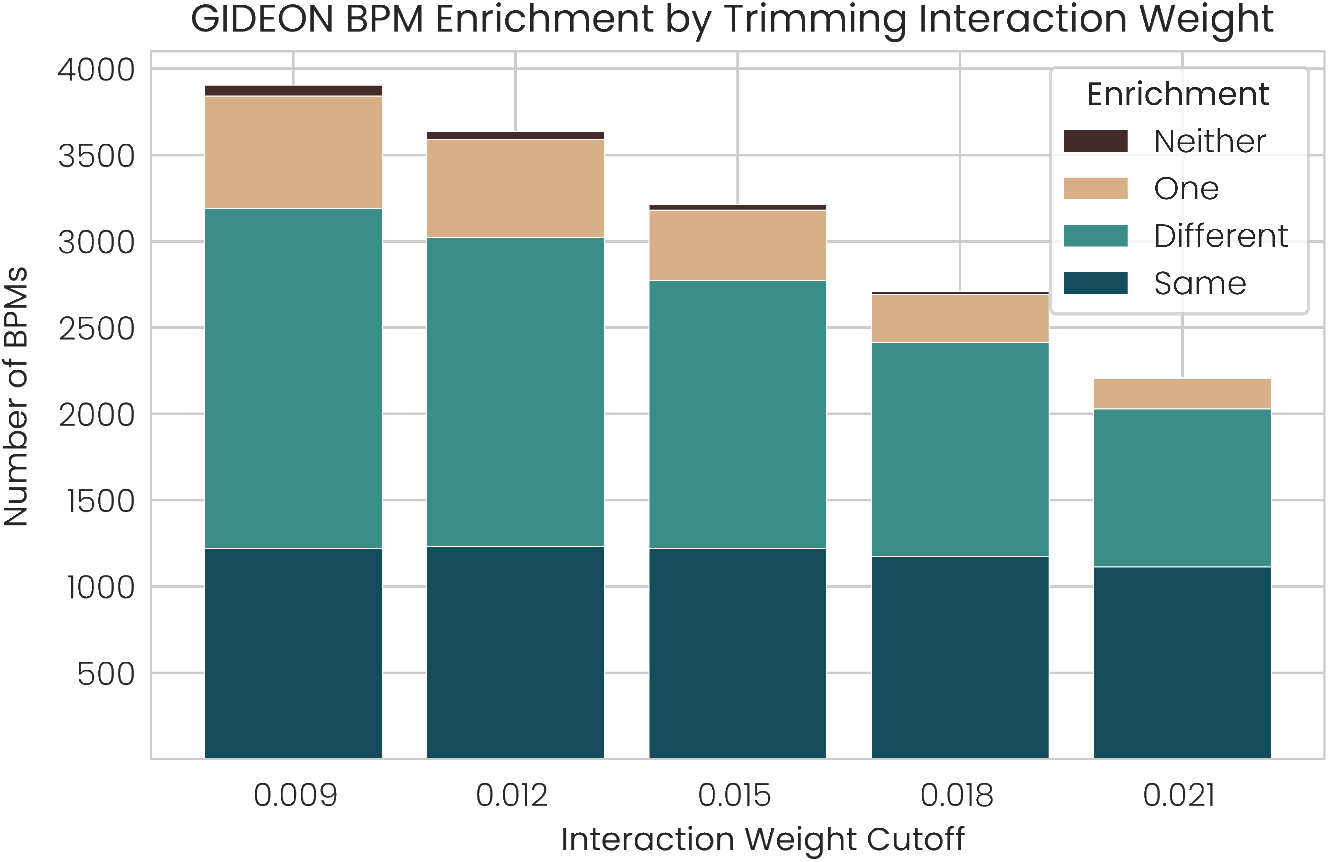
When varying the interaction weight cutoff for the trimming procedure of GIDEON with our DI weights, we identified that a cutoff of 0.015 produced cohesive BPMs itself, with surrounding cutoffs producing cohesive BPMs as well.

**Figure B.2:**
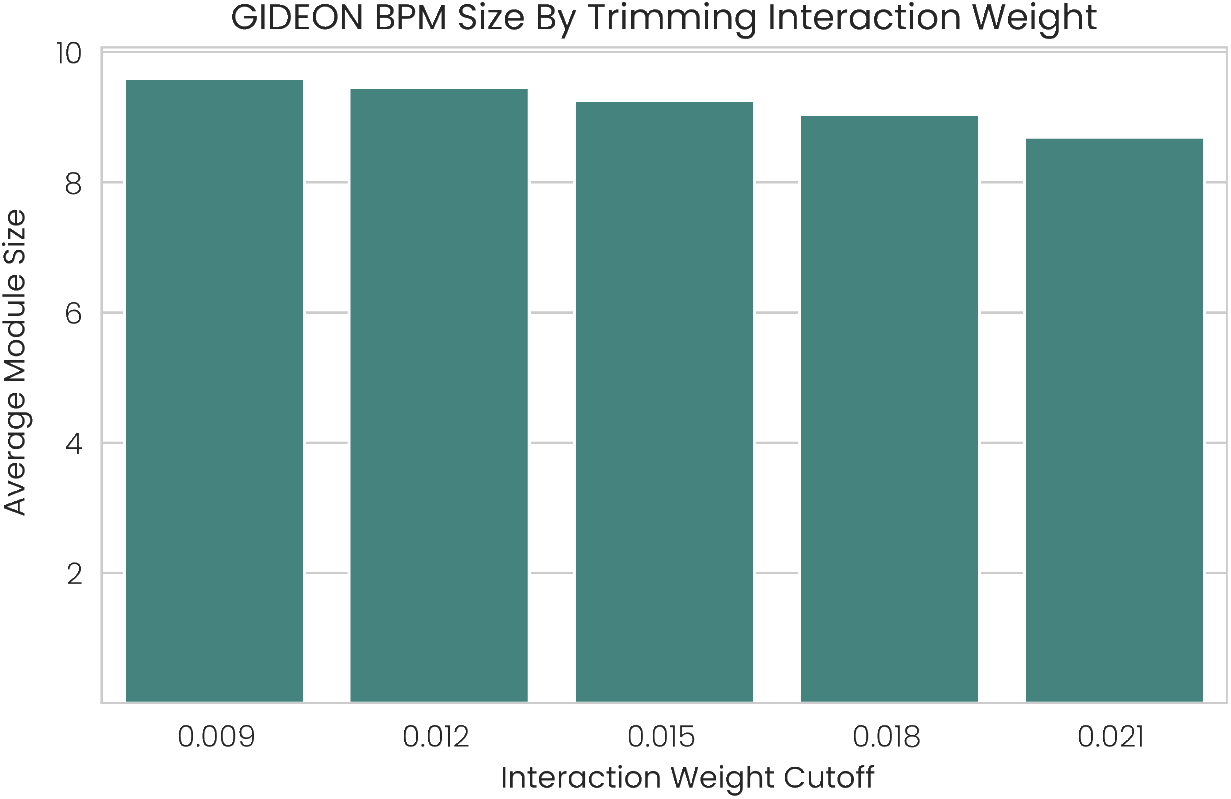
In addition to robust enrichment with a cutoff of 0.015 in Figure B.1, we find that varying the interaction weight cutoff has little impact on the size of outputted BPM modules, with the average number of genes only slightly dropping as the cutoff becomes stricter.

**Figure B.3:**
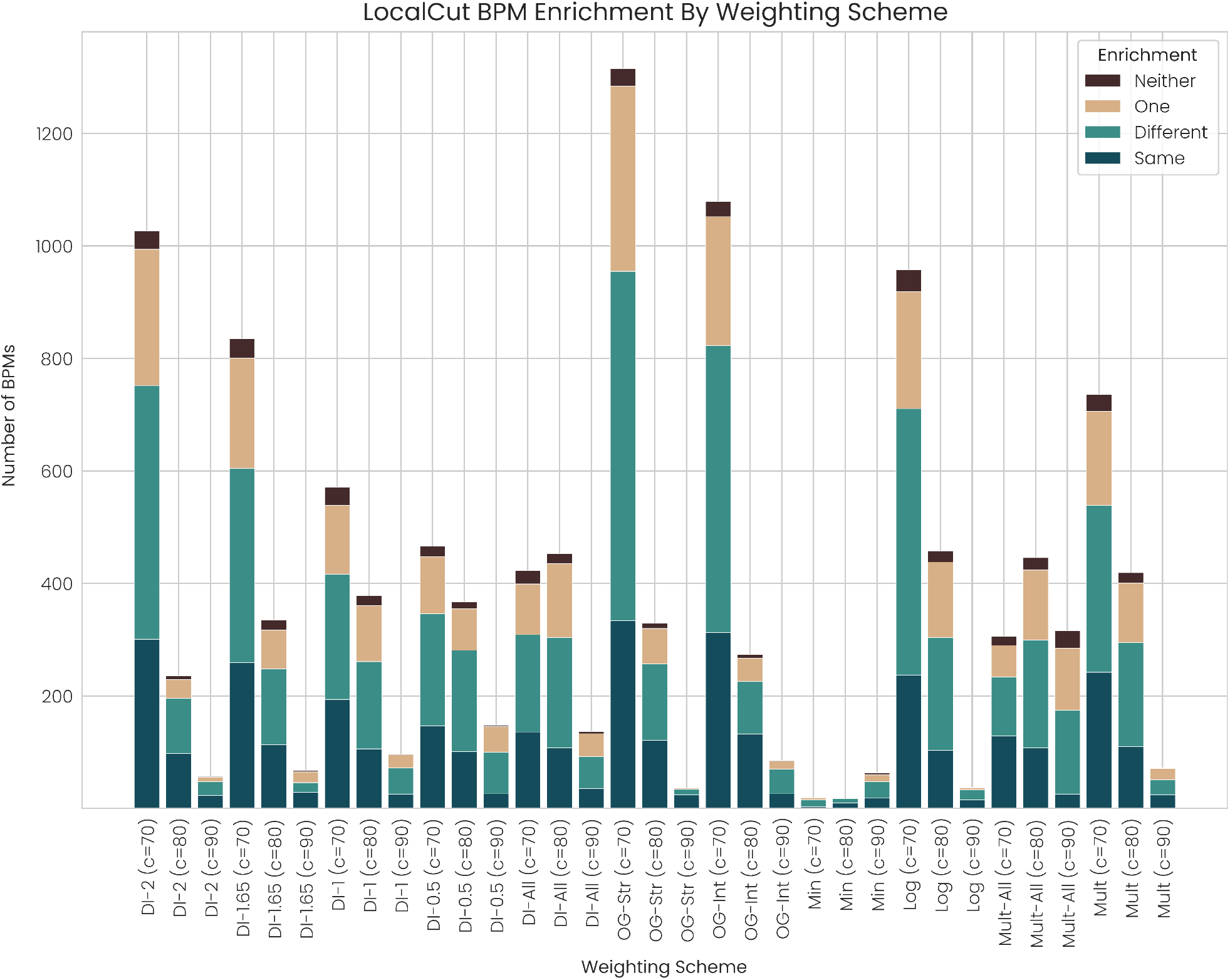
A comparison of LocalCut BPM output quantity and enrichment with different weighting schemes, with three settings of LocalCut’s *c* parameter. OG-Str and OG-Int weights for LocalCut outperform DI weights for LocalCut, but we again note that these methods require additional experimental data and are not generalizable to genetic interaction data. Note that LocalCut with DI weights already outperforms all approaches from [19] compared in [18]. We also compare LocalCut performance with our DI weights at varying *s*_*res*_ cutoffs, demonstrating that GIDEON’s default value of *s*_*res*_ *>* 2 produces also the most BPMs with competitive enrichment for LocalCut. Thus we set *s*_*res*_ identically for GIDEON and LocalCut when computing our DI weights for the results in the main paper.

**Figure B.4:**
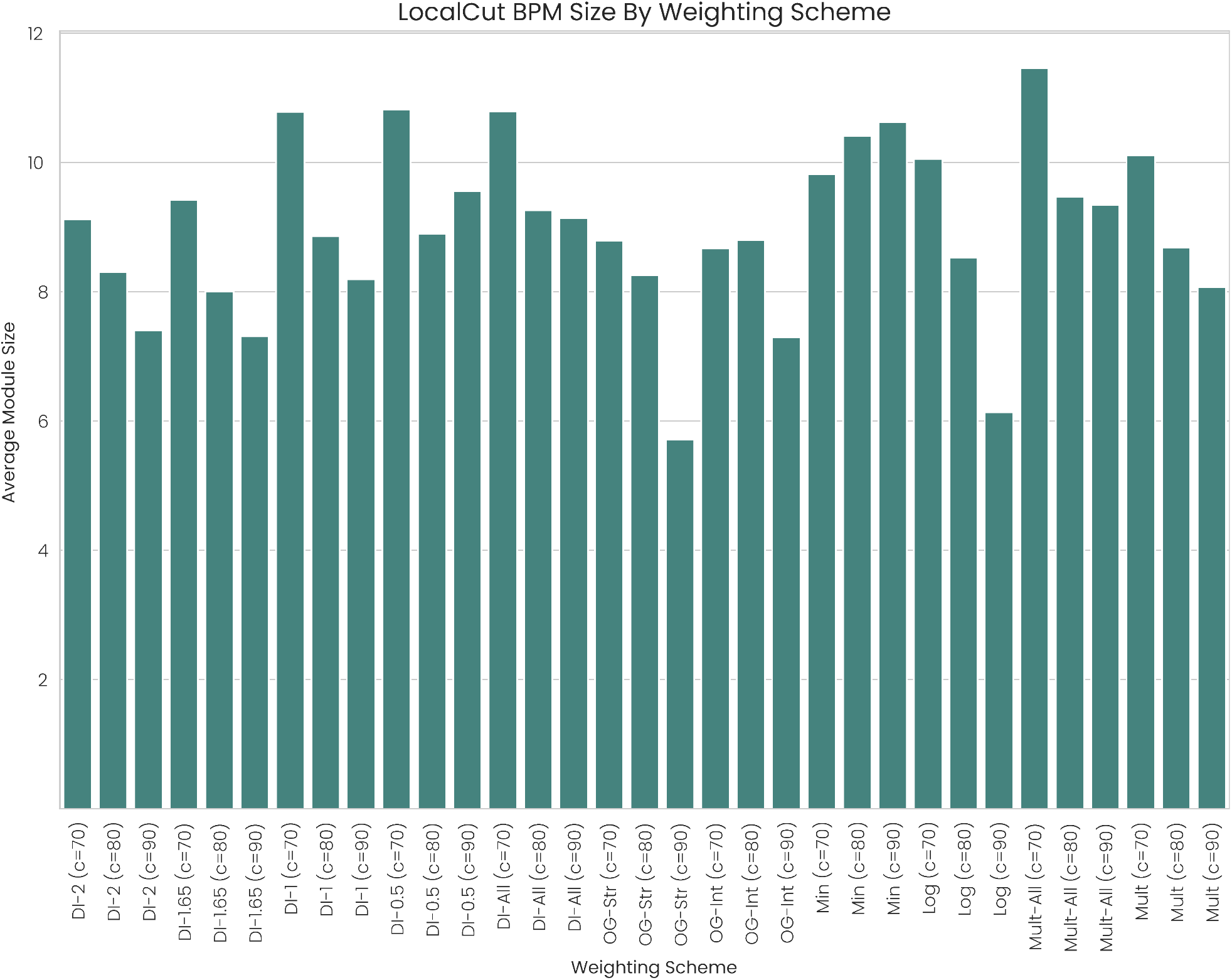
When considering LocalCut BPM output, we tend to favor weighting schemes and LocalCut parameters that produce slightly larger BPMs because they tend to yield more interesting biological insights. We note that our DI weighting scheme had an average module size of approximately 9 genes, the largest of the weighting schemes that produced a notable number of BPMs besides Log-All.

**Figure B.5:**
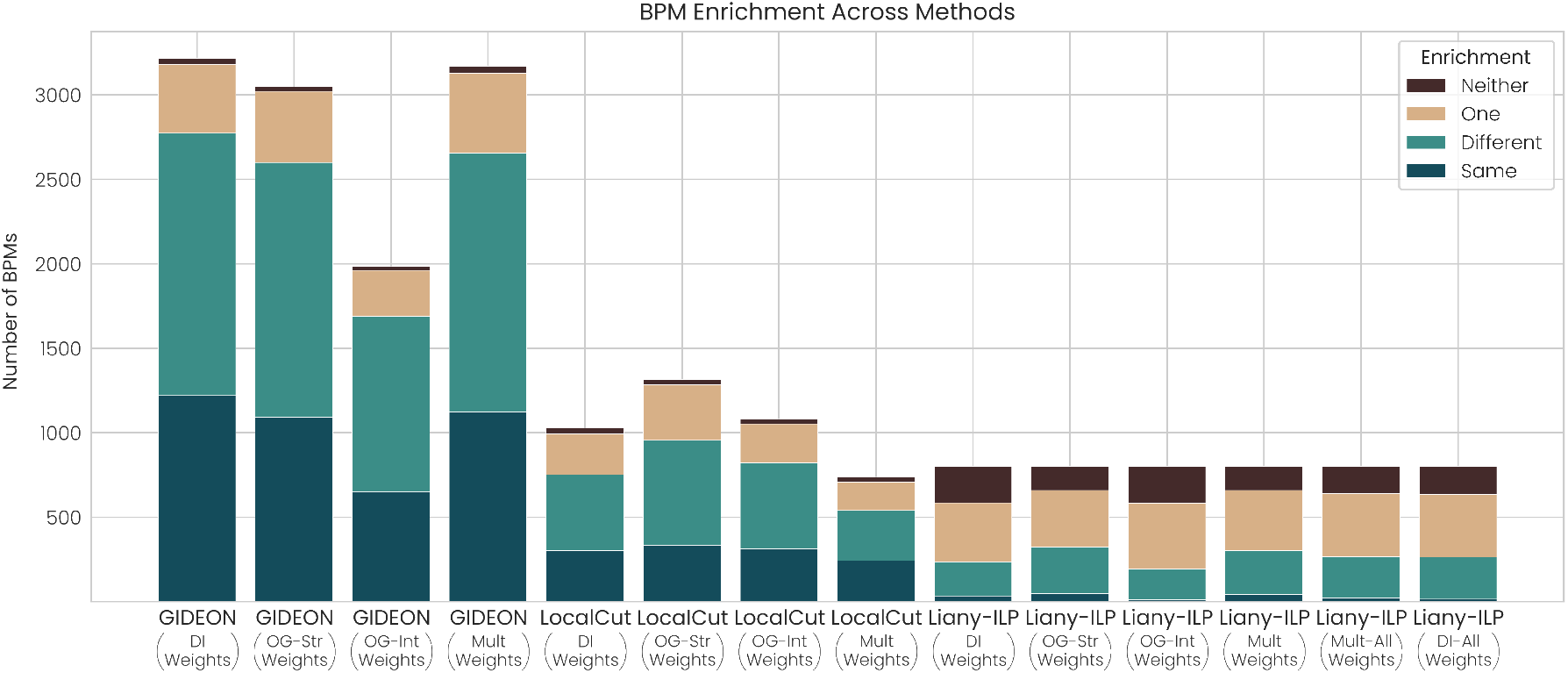
GIDEON substantially outperformed LocalCut and Liany-ILP both within and across weighting schemes. With GIDEON, our DI weighting scheme produced the most BPMs and the strongest enrichment scores, with OG-Str and Mult producing competitive BPM counts and enrichment as well. With LocalCut, the OG-Str weights identified 288 more BPMs than the DI weights, but we reiterate that additional experimental data is needed to build the OG-Str and OG-Int networks. With Liany-ILP, the OG-Str and Mult weights identified slightly more BPMs enriched for the same and different terms, but the poor enrichment scores of Liany-ILP make us hesitant to view these results as reflective of weighting scheme quality. Given that our DI weighting scheme outperformed OG-Sig with GIDEON and the results on LocalCut and Liany-ILP are competitive, our new approach is robust across methods, suggesting its potential for robust results across applications of epistasis data as well.

**Figure B.6:**
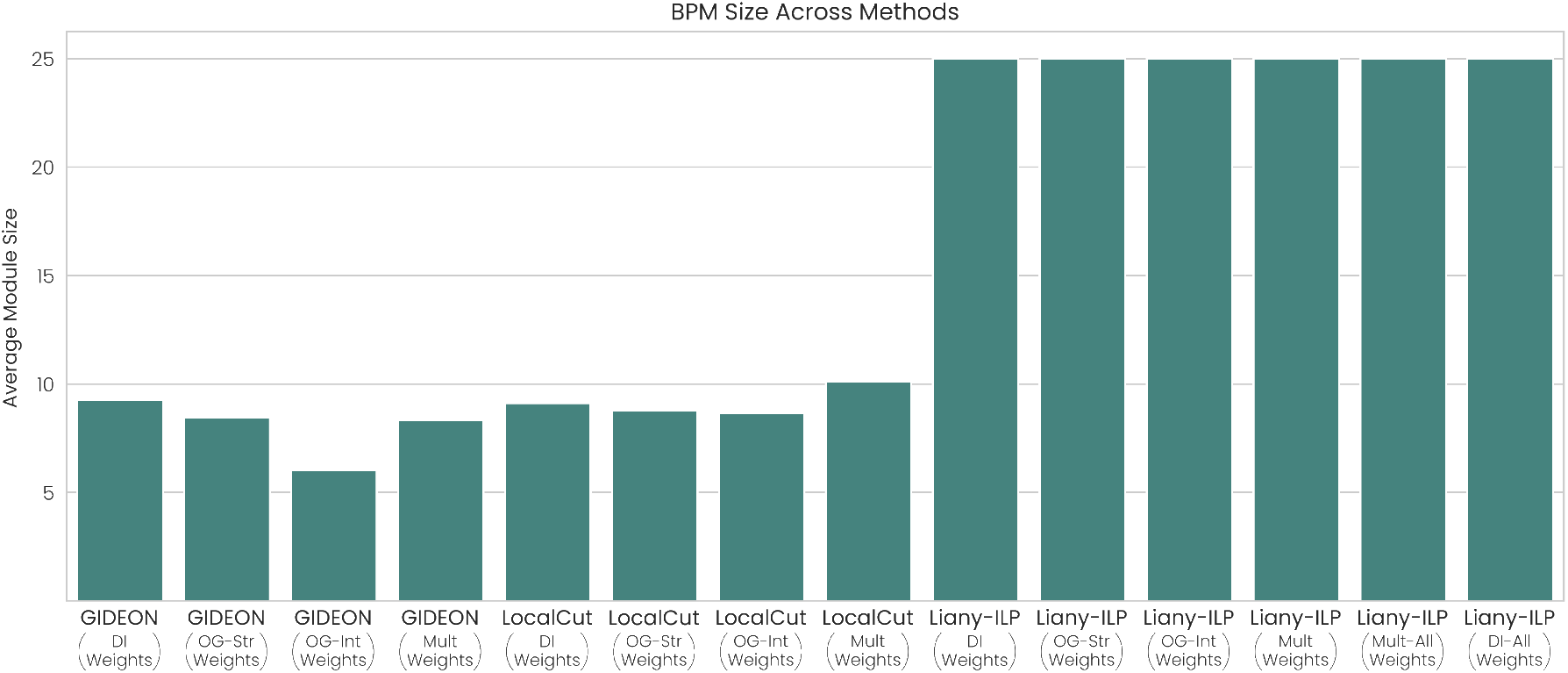
While GIDEON and LocalCut maintainted average BPM sizes of roughly nine genes in each module, Liany-ILP struggled with almost always producing BPMs with the maximum number of genes in each module. Initially exhibited by GIDEON and motivating the trimming procedure, the decision in Liany-ILP to ignore edges within modules exacerbated the problem of many genes in the network at least minimally improving the objective function, leading to the large BPMs.

**Figure B.7:**
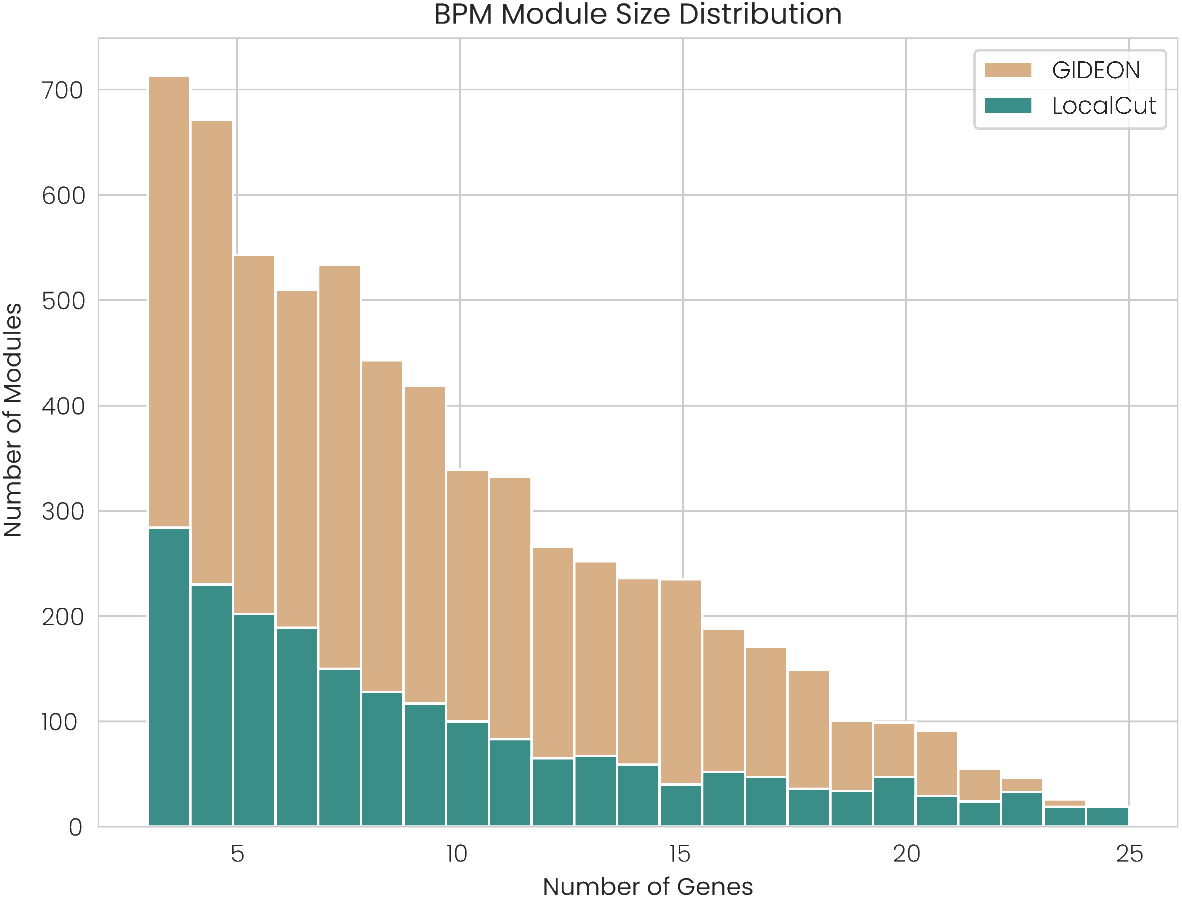
With our DI weights, GIDEON returns BPMs with the same average module size as LocalCut, demonstrating that two stages of pruning produce BPMs of reasonable size from the original ILP outputs with 25 genes in each module. Liany-ILP is not included because nearly every module had 25 genes.

**Figure B.8:**
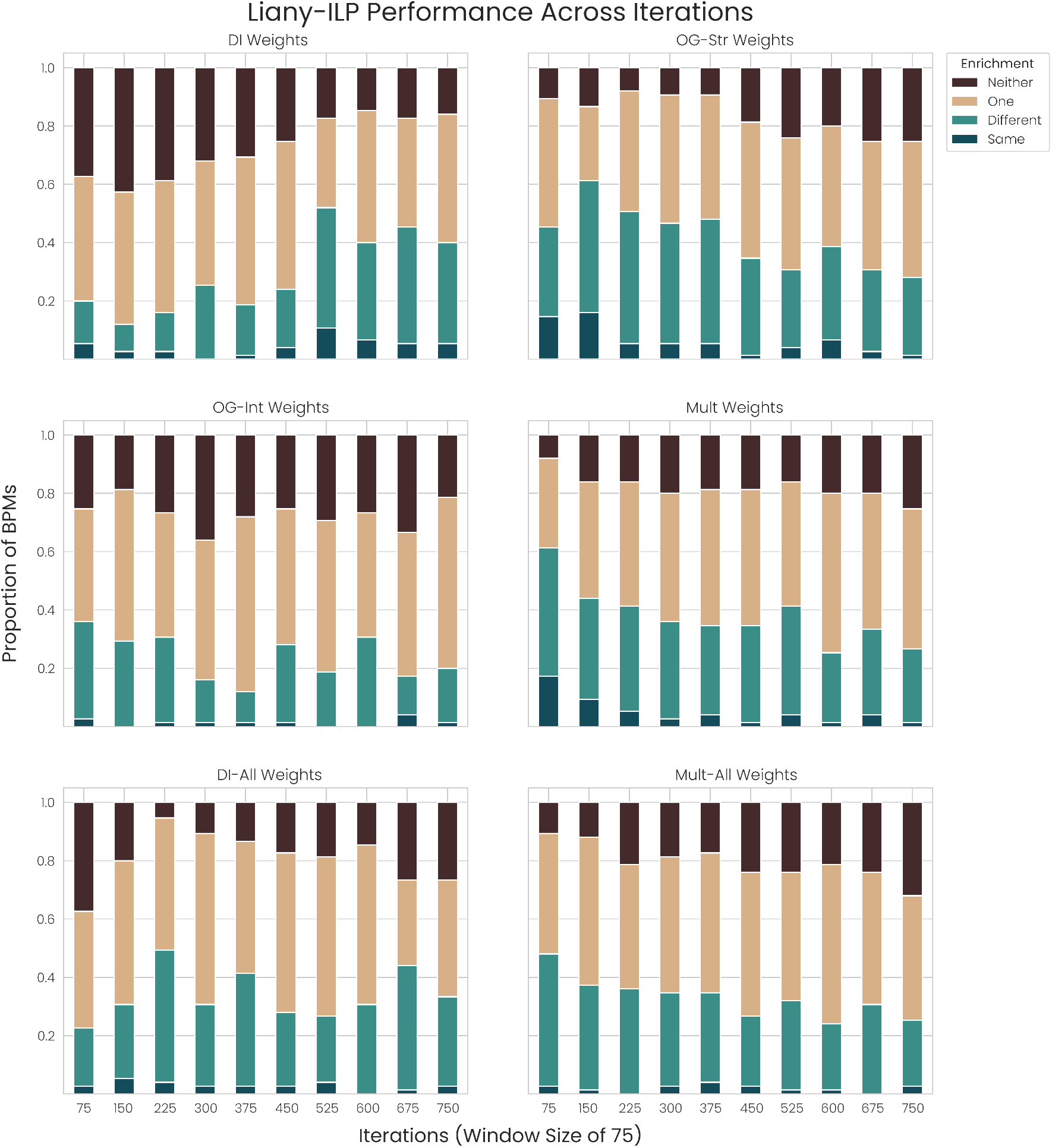
Given the sequential and iterative nature of Liany-ILP, the BPMs identified become worse as the algorithm continues. In other words, the first BPMs will tend to be best and BPM quality will fall as *K* (the number of BPMs) increases because Liany-ILP removes all edges after finding a BPM. We observe above that the BPM quality of Liany-ILP is poor from the first 75 BPMs and tends to fall from then on, demonstrating that higher values of *K* are unlikely to improve output. Because the edge removal is an inherent limit on the number of BPMs found, we also run Liany-ILP on Mult-All and DI-All but do not observe improvement in BPM quality. We also not that running Liany-ILP with the DI weights until the graph was empty produced 618 additional BPMs with only 18 enriched for the same term, further demonstrating that a larger *K* is almost certainly not beneficial. The sequential nature of Liany-ILP also lends itself to tedious runtimes, with identifying 833 BPMs using the Mult weights requiring over two days.

**Figure B.9:**
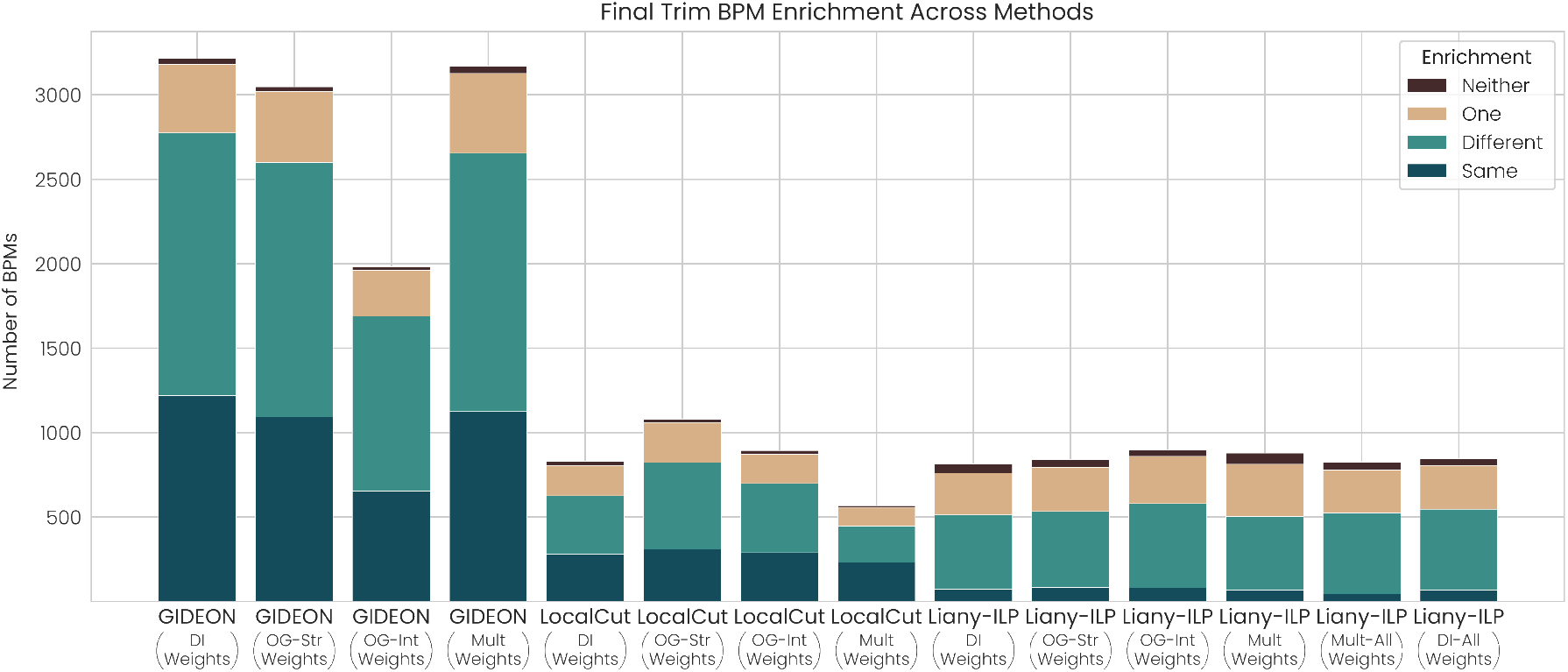
Finally, we compared GIDEON to new versions of LocalCut and Liany-ILP having these methods also apply the identical Final Trim step as GIDEON. Note that LocalCut loses BPMs through the Final Trim, demonstrating its prior performance is largerly dependent on disconnected BPMs. Liany-ILP performance slightly improves, though BPM enrichment remains poor.

**Figure B.10:**
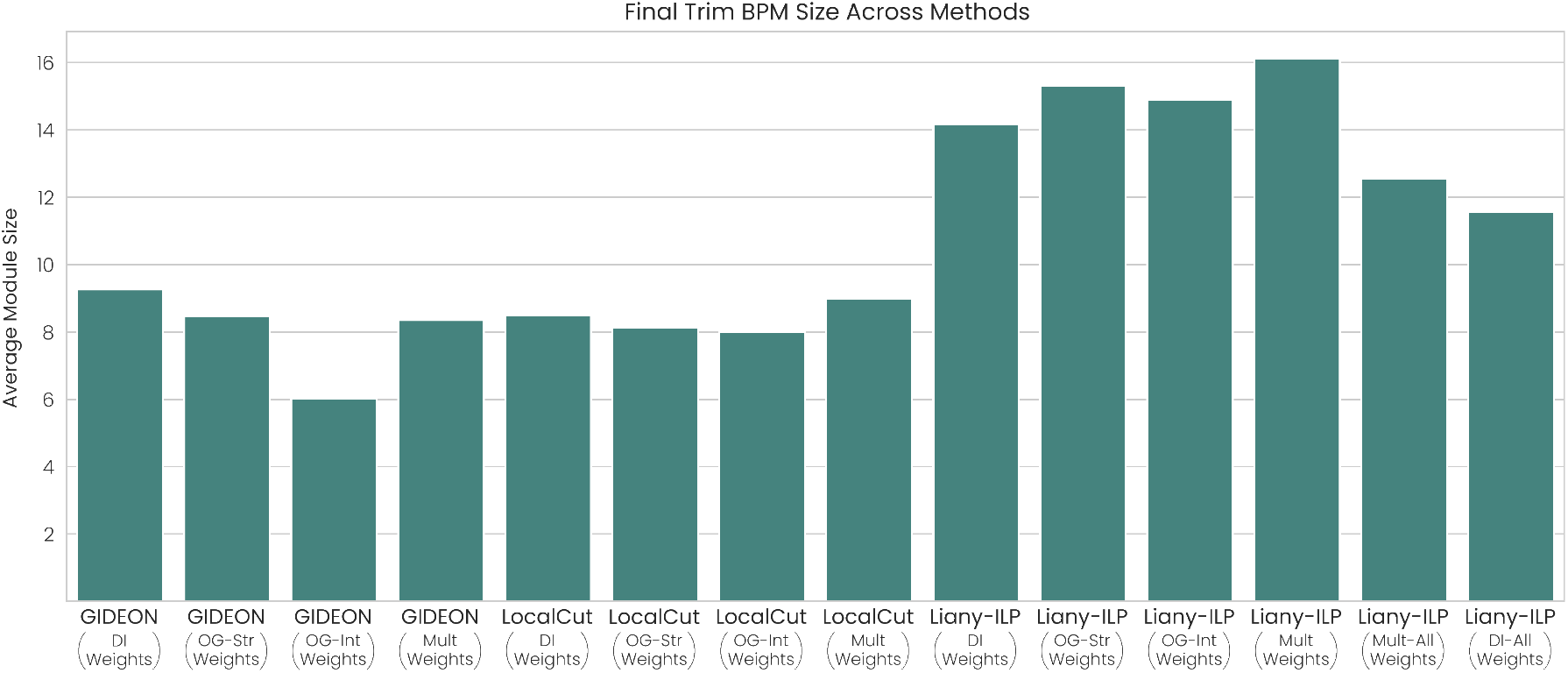
Compared with LocalCut and Liany-ILP after the Final Trim, GIDEON and LocalCut see an average of 8-9 genes per BPM than before while Liany-ILP begins to exhibit reasonable BPM sizes of 14-16 genes per BPM.

